# A confirmatory, dual-centric non-human primate study on the efficacy of novel oropharyngeal spray immunization with an adenoviral vector vaccine against RSV – Important lessons learned

**DOI:** 10.64898/2026.04.16.718916

**Authors:** Matthias Tenbusch, Gerrit Koopman, Petra Mooij, Berit Roshani, Pascal Irrgang, Dennis Lapuente, Ivanela Kondova, Willy M. Bogers, Edmond J. Remarque, Ramona Vestweber, Samuel Alberto Merida Ruiz, Nadine Krüger, Sebastian Meyer, Olaf Gefeller, Christiane Stahl-Hennig, Klaus Überla

**Affiliations:** Harald zur Hausen Institute of Virology, Uniklinikum Erlangen, Friedrich-Alexander-Universität Erlangen-Nürnberg (FAU), Erlangen, Germany; FAU Profile Center Immunomedicine, Friedrich-Alexander-Universität Erlangen-Nürnberg, Erlangen, Germany; Biomedical Primate Research Centre, Rijswijk, Netherlands; German Primate Center, Göttingen, Germany; Department of Medical Informatics, Biometry and Epidemiology, Friedrich-Alexander-Universität Erlangen-Nürnberg, Erlangen, Germany

## Abstract

In a confirmatory study, we evaluated the immunogenicity and protective efficacy of a heterologous prime-boost vaccination strategy against respiratory syncytial virus (RSV) in non-human primates. Building on prior evidence of protective mucosal immunity induced by intramuscular DNA priming followed by an oropharyngeal adenoviral boost, we conducted a randomized, blinded, dual-centre study across two European primate research facilities. Rhesus macaques received a codon-optimized RSV-F DNA vaccine via electroporation, followed by two mucosal administrations of a recombinant adenovirus serotype 5 vector encoding the same antigen. Control groups included animals vaccinated with irrelevant influenza antigens and a comparator group mimicking natural immunity induced by primary RSV infection.

Systemic and mucosal immune responses, including RSV-F-specific antibodies and tissue-resident memory T cells, were monitored longitudinally. Here, we detected robust immune responses, but with some variability between the two centres. However, following experimental RSV challenge performed 22 weeks after the final immunization, RSV-vaccinated animals demonstrated markedly reduced viral replication in both upper and lower respiratory tracts. However, unexpected RSV-specific immunity in the control group at one single study site prevented confirmation of the predefined primary endpoint.

Overall, these results support the potential of mucosal adenoviral boosting following DNA priming to induce protective immunity against RSV, while highlighting challenges associated with multi-centre preclinical vaccine studies.

## Introduction

Respiratory syncytial virus (RSV) remains a major cause of acute lower respiratory tract infection, producing substantial morbidity and mortality particularly among infants, young children and older adults. Nearly all children experience an infection within their first 2 years with the incidence peaking in the first three months of life^1,2^. Primary infection compromises the lower airway in 15-50% of cases requiring in-patient treatment in 1-3% ^3^. Globally, it was estimated that annually about 30 million of RSV-induced episodes of acute lower respiratory tract infection occurred leading to about 60.000 deaths in children younger than five years of age^4^. In elderly or immunocompromised individuals, the infection rates and disease severity are comparable to seasonal influenza infections. Since natural infections do not induce persistent immunity^5–7^, the development of vaccines providing efficient and long-lasting protection against RSV were a major topic in the field. Until 2022, the only preventive measure was a passive immunization with an RSV-F specific monoclonal antibody (Palivizumab), which was administered to high-risk groups of children only. Currently, many countries recommend a passive immunization of all new-borns before their first RSV season with Nirsevimab, a newly designed, half-life extended mAb against RSV-F. In clinical trials, Nirsevimab markedly reduced medically attended RSV disease and severe disease outcome^8–10^. In 2023, the FDA approved the two protein vaccines based on the stabilized prefusion RSV-F protein, Arexvy (GlaxoSmithKline) and ABRYSVO^TM^ (Pfizer), for prevention of RSV in the elderly. Both vaccines demonstrated efficacy against severe lower respiratory tract infection in clinical phase III trials ^11,12^. In addition, ABRYSVO^TM^ is licensed for the use as maternal immunization of pregnant women to protect new-borns in the first months after birth^13^. Most recently, a mRNA vaccine produced by Moderna complements the current portfolio of RSV prophylaxis measures^14,15^.

Despite these improvements, passive immunization and intramuscularly applied vaccines do not produce durable mucosal immunity in the upper respiratory tract, which is the primary site of entry and early replication of RSV and of most other respiratory viruses. Thereby, these preventive measures can reduce severe disease, but might have limited impact on viral shedding and onward transmission as witnessed during the recent COVID19 pandemic by the large number of SARS-CoV-2 breakthrough infections in fully vaccinated individuals^16–18^.

Therefore, huge affords were made in developing new vaccine strategies focusing on mucosal immunity to limit respiratory tract infections by RSV, Influenza A or SARS-CoV-2^18–25^. In case of RSV, efficacy of mucosal vaccines have been reported in several preclinical animal models or even in human challenge studies^19,23,26–28^.

We demonstrated that an intramuscular DNA immunization followed by an oropharyngeal spray application of an adenoviral vector induced long-term mucosal immunity to RSV-F in non-human primates (NHP)^29^. In this study, RSV-F specific IgA antibodies and F-reactive T-cells were detected in bronchoalveolar lavage up to ∼100 weeks after the mucosal booster immunization. After an additional DNA immunization at that time, vaccinated rhesus macaques exhibited significantly reduced viral loads in the upper and lower respiratory tract after an experimental RSV challenge infection. However, the late DNA boost did not impact the mucosal immune response and most probably did not contribute significantly to the observed protection. To demonstrate the protective capacity of the DNA prime-adenoviral vector boost regimen, we performed a confirmatory study with a more condensed and stringent immunization schedule consisting of one intramuscular DNA application followed by two mucosal rAd immunization and a final RSV challenge 22 weeks after the last immunization. Furthermore, we intended to increase the external validity of our study by performing a randomized, blinded non-human primate study at two European primate centres. To the best of our knowledge, such a harmonized study on immunogenicity and efficacy of an experimental vaccine has never been done before.

At each centre, we had three treatment groups with eight animals per group at the German Primate Centre (DPZ) and four animals at the Dutch Biomedical Primate Research Centre (BPRC). Our vaccine group initially received an DNA vaccine encoding a codon-optimized RSV-F via electroporation and then two oropharyngeal spray immunizations with Ad5-F, encoding the same antigen. As a control group, mock-treated animals received the same types of vaccines encoding irrelevant influenza antigens. Finally, a third comparator group was included to mimic natural immunity established by primary RSV infection, which does not prevent re-infections in humans. We analysed systemic and mucosal antibody and tissue-resident memory T-cell (TRM) responses longitudinally and confirmed the vaccineś immunogenicity with centre-specific differences. Furthermore, the RSV-immunized animals showed very low levels of viral replication after the experimental RSV challenge infection in both centres. However, due to some unexpected findings in the control group at the DPZ, the confirmatory endpoint was not reached in this multi-centric study.

## Methods

### Study design

This study was performed at the German Primate Center (DPZ, Göttingen, Germany) and the Biomedical Primate Research Center (BPRC) in Rijswijk, The Netherlands. All treatments and sampling procedures were performed after intensive discussion among consortium partners and sharing of common protocols, but in a completely independent manner to ensure external validity of our demonstrated vaccine efficacy. A pre-registration of the study with the title “RSV-Protect” was done at animalstudyregistry.org.

#### Sample size calculation

Due to data from our previous study^29^, we knew that viral load data in vaccinated and unvaccinated groups were skewed and showed strong heterogeneity in variability between groups. Therefore, standard statistical procedures assuming normality for sample size calculation and data analysis could not be applied. For calculating sample sizes, we simulated Weibull distributions parameterized to capture discrepancy, skewness, and heterogeneity in variability from the previous data. We then drew random samples of equal size per group from these simulated distributions in a Monte Carlo experiment to compute the statistical power of the Mann-Whitney(-Wilcoxon) test with a two-sided significance level of 0.05 over a range of 36 different distributional scenarios and varying sample sizes. A sample size of at least 12 per group yielded a power of > 0.90 in scenarios compatible with assumptions of effect size backed by the previous data.

#### Methods for randomization

To reduce the risk of bias, the monkeys used in the study were randomly assigned by the statistician to the three groups. Randomization was stratified for sex, weight and age. When animals were already pair-housed, randomisation was performed pairwise.

#### Methods of blinding

Based on the randomization results, the vaccine producer filled the vaccines in vials labelled only with the designation of the respective recipient monkeys and the vaccine time point. The staff performing the immunizations and the RSV challenge at DPZ and BPRC and the staff performing viral load determinations and immune monitoring were blinded for the RSV and Mock groups, since only the statistician and the person preparing the vaccine batches knew the group assignment. Due to different vaccination routes and time schedules the comparator group could not be injected in a blinded manner, but lab workers were kept blinded by labelling samples with monkey designation, sample type and time point only.

### Animals, housing and ethics statements

The study was performed with a total number of 36 adult rhesus macaques (*Macacca mulatta*). At the BPRC, 12 female rhesus macaques at the age between 4-13 years were included, while at the DPZ six female and 18 male rhesus macaques at an age between 4-10 years were treated. The distribution and specifications of the animals per group at each center are shown in Table 1.

**Table 1:** Overview on sex, age, and weight distribution of test animals at the different centers*.

|  | Mock |  | RSV |  | Comparator |  |
| --- | --- | --- | --- | --- | --- | --- |
|  | DPZ | BPRC | DPZ | BPRC | DPZ | BPRC |
| Sex;<br>(no. f/no. m) | 2/6 | 4/0 | 2/6 | 4/0 | 2/6 | 4/0 |
| Age;<br>median (ICR)<br>[range] | 7.5<br>(7-8.75)<br>[4-9] | 8.0<br>(7.25-11)<br>[7-12] | 8.0<br>(6.25-9)<br>[5-10] | 6.5<br>(4-12)<br>[4-13] | 7.5<br>(5.5-8)<br>[4-8] | 7.0<br>(6-11.75)<br>[6-13] |
| weight;<br>median (ICR)<br>[range] | 11.8<br>(8.2-13.5)<br>[7.4-14.4] | 6.9<br>(6.7-9.0)<br>[6.7-9.7] | 9.7<br>(7.5-13.6)<br>[6.1-15.5] | 7.8<br>(6.1-8.9)<br>[5.6-9.2] | 10.5<br>(5.9-12.8)<br>[5-16.4] | 8.2<br>(6.1-11.4)<br>[6-11.9] |

\*no, number; f, female; m, male; ICR, Interquartile Range

Before inclusion, all animals underwent a full physical examination and clinical chemistry and hematology evaluation. Only healthy individuals with normal clinical chemistry and hematology levels and without pathogens were selected.

All housing and animal care procedures at the BPRC in Rijswijk, the Netherlands, are in compliance with European directive 2010/63/EU, as well as the Standard for Humane Care and Use of Laboratory Animals by Foreign Institutions provided by the Department of Health and Human Services of the United States National Institutes of Health (NIH, identification number A5539-01). The BPRC is accredited by the American Association for Accreditation of Laboratory Animal Care (AAALAC). The study was reviewed and approved by the Dutch “Centrale Commissie Dierproeven” (AVD50200202114508 and AVD50200202114680) according to Dutch law, article 10a of the “Wet op de Dierproeven”, and BPRC’s Animal Welfare Body (IvD).

Animals housed at the German Primate Center (DPZ) were cared for by experienced animal caretakers according to the German Animal Welfare Act complying with the European Union guidelines on the use of non-human primates for biomedical research and the Weatherall report. The animal experimentations were approved by the Lower Saxony State Office for Consumer Protection and Food Safety under the project license AZ. 33.19-42502-04-21/3752. In agreement with §11 of the German Animal Welfare act, the DPZ has the permission to breed and house non-human primates under the license 392001/7 granted by the local veterinary office.

Animals were housed at the BPRC in ABSL-3 facilities (building OG at the BPRC) and ABSL-3** at the DPZ. During the experiment the animals were pair-housed if socially compatible. In case of incompatibilities between animals, which could have been detrimental to the animal welfare, animals were kept in individual cages, but with constant visual, acoustic and olfactory contact to their neighbours. In both centres the light schedule is 12 hours of light and 12 hours of dark controlled centrally. The monkeys were offered a daily diet consisting of monkey food pellets (BPRC: Hope Farms, Woerden, The Netherlands, DPZ: Safe 307 Primates extruded Diet, SAFE, Augy, France; Altromin 6029, Altromin Spezialfutter, Lage, Germany; Ssniff V3986-000, Sniff Spezialdiäten, Soest, Germany), fruit and vegetables. Enrichment (toys, extra food) was offered daily. Drinking water was available ad libitum.

### Vaccines and RSV virus

The DNA vaccines are all based on the pVAX plasmid encoding codon-optimized sequences for RSV-F, IAV-HA or IAV-NP (both derived from H1N1/Puerto Rico/8/34) and were described before ^30,31^. The adenoviral vectors (AdV) encoding the same antigens were reproted as well^32,33^

. High-titer stocks were generated and purified by SIRION Biotech (Martinsried, Germany). The vaccine doses of 2 mg DNA/dose or 2×10^9^ infectious units (IU) Ad5/dose in sterile NaCl 0,9% were prepared and labeled at the virology laboratory in Erlangen and then sent to the two centers in a blinded manner.

For the primary and challenge infection, macaque-adapted RSV^34^ was propagated on TMK cells and subsequently aliquotted and stored at −135°C. The titer of this challenge stock as measured on HEp-2 cells was 1 × 10^5^ PFU/ml.

### Immunizations, treatments and sampling

For all experimental procedures performed at DPZ, rhesus macaques were anesthetized by i.m. injection of a mixture of 5 mg ketamine, 1 mg xylazine and 0.01 mg atropine per kg body weight. At the BPRC animals were anesthetized with a mixture of 5 mg ketamine and 0.05 mg medetomidine per kg body weight and immediately after the procedure atipamezole was given for faster recovery (0.25 mg per kg). Blood samples were collected from the femoral vein using the vacutainer system (BD). Immunization strategy included a prime immunization with DNA vaccines via intramuscular electroporation followed by two booster immunizations using a spray applied to the tonsils at weeks 8 and 28 (Figure 1). For DNA vaccination, 2 mg DNA was diluted in isotonic 0.9% saline solution and delivered into both quadriceps muscles using the TriGrid Delivery System following the manufacturer’s protocol (Ichor Medical Systems, San Diego, CA, USA). Booster immunizations were performed by applying 2×10^9^ IU Ad5 diluted in isotonic 0.9% saline solution to both tonsils using MADgic Laryngo-Tracheal Mucosal Atomization Device (MAD720, Teleflex, Wayne, PA, USA). The comparator group was challenged intranasally with RSV (10^5^ PFU) at week 8 followed by two immunization with seasonal quadrivalent influenza vaccine (Influvac Tetra Saison 2021/2022) at weeks 24 and 28. At week 40 rhesus macaques from all groups were inoculated intranasally with 10^5^ PFU RSV.

**Fig. 1:**
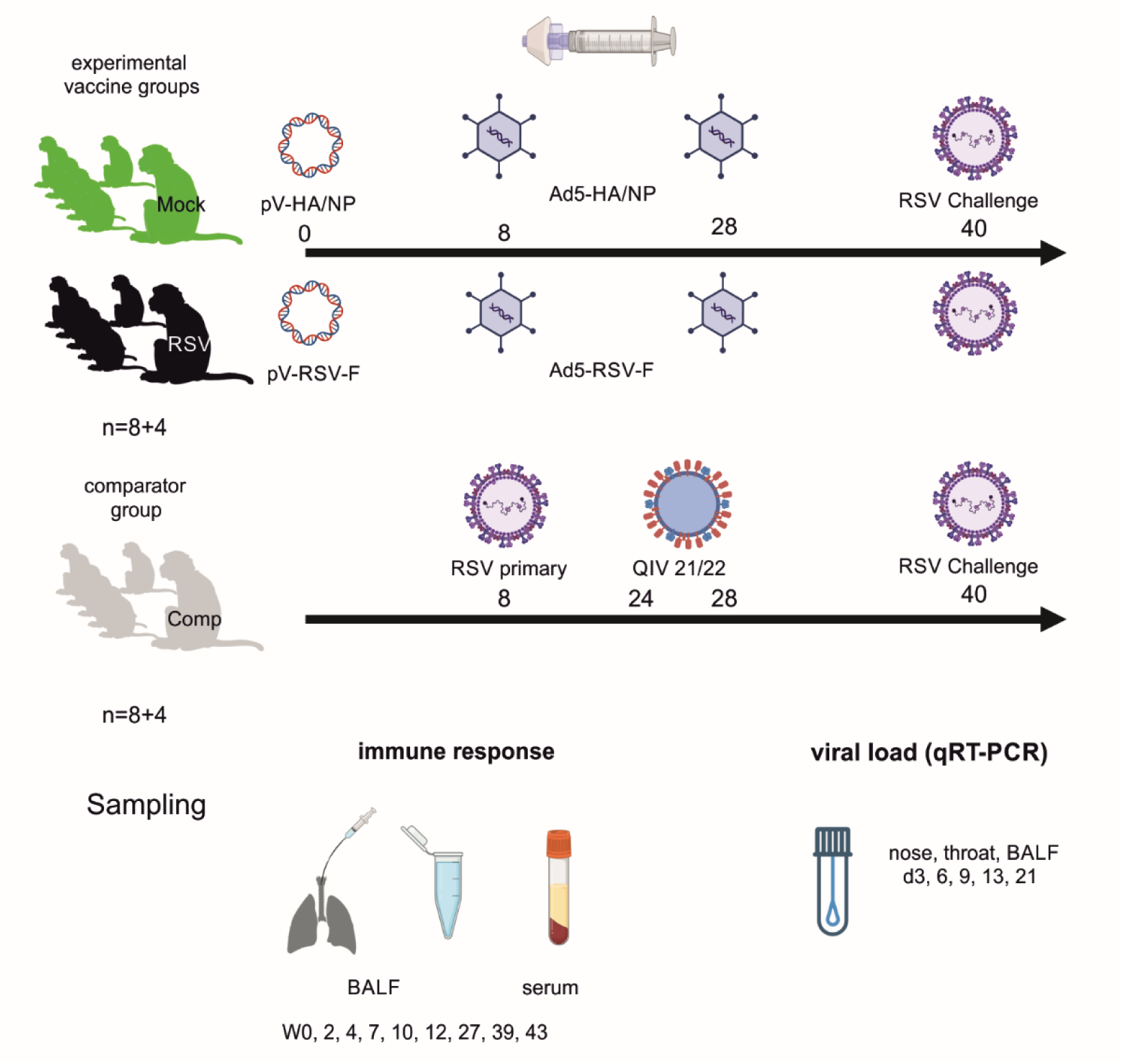
Experimental schedule. In the two experimental vaccine groups, rhesus macaques were immunized via intramuscular DNA electroporation with 2mg of plasmid DNA encoding either RSV-F or IAV-HA/NP as antigens. At week 8 and week 28, these animals were boosted via oropharyngeal spray application of 2×10^9^ IU Ad5 vectors encoding the same respective antigens. The comparator group were exposed to 1x 10^5^ PFU of RSV in week 8 and received two doses of inactivated influenza vaccines (QIV) at week 24 and 28. To evaluate vaccine efficacy, all animals were exposed to 1x 10^5^ PFU of RSV at week 40 and viral load was monitored. BALF and sera for analyses of humoral and cellular immune responses were collected at the indicated time points, as well as nose and throat samples and BALF for the detection of viral RNA after RSV exposure. Created in BioRender. Tenbusch, M. (2026) https://BioRender.com/80hpbbl

#### Nasal and oropharyngeal swabs

Nasal sampling was carried out by gently swabbing each nostril with a Minitip flocked swab (80 mm, Copan®). Next, swabs were placed together into a single collection tube containing 1 mL of viral transport medium (VTM). The oropharynx was sampled using a single regular flocked swab (80 mm, Copan®), which was gently swept across the posterior pharyngeal wall. Thereafter, the swab was placed into a collection tube containing 1 mL of viral transport medium (VTM). All samples were kept on ice until further processing.

#### BAL collection

BAL cells were obtained by placing a flexible bronchoscope (4 EB-530P Video bronchoscope, Øbiopsy channel 1,2mm, L=600mm; Fujifilm Germany, Ratingen) into the trachea to the bifurcation of the main bronchi and then into one of the main bronchi until the wedge position was reached. For determination of viral load after RSV exposure, a small volume of BAL fluid (BALF) was collected. A bolus of 5-8 ml (DPZ) or 3-5 ml (BPRC) of body-warm 0.9% saline solution (Ecolav NaCL 0.9%, B.Braun) was administered through the working channel using a

5 or 10 ml syringe (Omnifix solo, B.Braun). The channel was immediately flushed with 1ml air to push through most of the dead volume of the fluid retained in the channel. Then, the fluid was gently aspirated. Recovered fluid was kept on ice and later stored in aliquots at -80° C. For isolation of BAL cells and antibody measurements, a total volume of 60 ml body-warm 0.9% saline solution (Ecolav NaCL 0.9%, B.Braun) was used. At DPZ, initially, a bolus of 30 ml was delivered through the working channel using a 30 ml syringe (BD Plastipak Luer-Lock syringes, BD). This syringe was then removed, and a 10 ml syringe (Omnifix solo, B.Braun) was connected to recover the fluid in portions by gentle aspiration. The procedure was repeated for the second bolus of 30 ml. At the BPRC, the volume of 60 mL was given in three portions of 20 mL, using a 20 mL syringe. After each 20 mL injection the fluid was recovered with the same syringe. Either the left or right side of the lung was treated.. Recovered fluid was stored on ice. Obtained BALF was passed through a 70µm cell strainer (Falcon™ Cell Strainers, Corning) and washed twice with PBS/5% fetal bovine serum (FBS, anprotec). Cells were counted in the presence of trypan blue (Sigma-Aldrich). A sample was considered as clean and easy to evaluate if it contained only large, granular lymphocytes. Following another centrifugation step, cells were resuspended in RPMI1640 (PAN-Biotech) with 10%FBS (anprotec) and 1%Penicillin-Streptomycin (PAN-Biotech) for further analysis.

### Intracellular cytokine staining of BAL cells

Antigen-specific cytokine responses of freshly isolated BAL cells were analyzed by intracellular cytokine staining (ICS). BAL cells were incubated with anti-CD28 (clone CD28.2, BioLegend, San Diego. CA, USA) and anti-CD49d (clone 9F10, BioLegend) antibody (each 1 µg/mL), and either a combination of phorbol-12-myristat-13-acetat (PMA) and ionomycin (both Sigma Aldrich, St.Louis, MO, USA: each 1 µg/mL), RSV F peptide pool (2 µg/mL/peptide) or medium only at 37°C for 6 hours in the presence of Brefeldin A (Brefeldin A solution, 1:1000, Biolegend). The RSV-F peptide pools consist of 138 overlapping peptides spanning the whole RSV-F protein (15-mers, overlapping by 11 AA, Genscript, NJ, USA). Cells were then washed with PBS and incubated with 50 µL of a mixture of a live/dead stain (Zombie Yellow Fixable Viability Kit, BioLegend) and antibodies for surface markers including CD3^AF700^ (clone SP34.2), CD4^V450^ (clone L200), CD45^PE^ (clone D058-1283, all BD Pharmingen), CD45RA^ECD^ (clone 2H4), CD103^FITC^ (clone 2G5,both Beckman Coulter, Brea, CA, USA), CD8a^PE-Cy5^ (clone RPA-T8), CD69^BV510^ (clone FN50), CD197^APC^ (clone G043H7) and CD279(PD-1)^PE-Cy7^ (clone EH12.2H7, all BioLegend) for 30 min at room temperature (RT) in the dark. Cells were fixed by adding RBC lysis/fixation solution (diluted 1:10 in dH2O, BioLegend) for 15 min at RT and in the dark. Subsequently, the cells were washed three times with intracellular staining permeabilization buffer (diluted 1:10 in dH2O, BioLegend), and resuspended in intracellular staining permeabilization buffer with anti-IL-2^BV650^ (clone MQ1-17H12, BioLegend), IL-10^AF647^ (clone JES3-9D7, BioLegend), anti-TNFα^BV711^ (clone Mab 11, BioLegend) and anti-IFNγ^PerCP-Cy5.5^ mAb (clone 4S.B3, BioLegend). After 20 min incubation at RT, cells were washed with the permeabilization buffer and resuspended in PBS/1%BSA. Acquisition was performed using spectral analyzers. At the BPRC the Cytek Aurora (Cytek, Fremont, CA, USA) and at DPZ the ID7000 (Sony Biotechnology, San Jose, CA, USA) were used. Analysis was performed using FlowJo software (Version 10.9, BD, Ashland, OR, USA). First, a lymphocyte gate was drawn. Then singlets were selected using a forward scatter-height (FSC-H)/forward scatter-area (FSC-A) plot, and CD3 positive cells that were negative for the live/dead marker were selected (gating strategy shown in Supplementary Fig. 2). Subsequently, total CD4 and CD8 as well as CD4 and CD8 TRM (CD45RA^-^ CCR7^-^ CD69^+^CD103^+^) subsets were analyzed for cytokine expression. Boolean gating was used on IFNg, TNFa, and IL-2 positive cells to analyze the Th1 cytokine expression patterns in detail.

### Flow-cytometric antibody assay

For the detection of RSV-F specific antibodies, doxycylin-inducible HEK293 cell lines expressing the respective antigen were used as target cells after 24h induction with 400ng/ml doxycycline as described before^19^. Briefly, 5×10^5^ HEK 293 cells were incubated for 20 minutes at 4°C with the respective biological sample (BAL or serum) diluted in 100 µl FACS-PBS (PBS with 0.5% BSA and 1 mM sodium azide) to bind to RSV-F on the surface. After washing with 200 µl buffer, antigen-specific antibodies were detected with cross-reactive anti-hIgG-FITC (jackson 109-096-088; 1:300) + anti hIgA-APC (abcam; ab99897; 1:200). After further washing, samples will be measured on a flow cytometer like the AttuneNxt (ThermoFisher) and analysed using FlowJo software (Tree Star Inc.). To quantify the RSV-F specific IgG antibodies, a standard curve was generated with defined concentrations of the monoclonal antibody palivizumab. For IgA levels in BAL, the mean fluorescence intensity (MFI) was directly shown.

### Neutralization assay

Neutralizing antibody titers were determined by using a recombinant RSV encoding for the firefly luciferase (RSV-Luc) in a 96-well neutralization format assay as described before ^35^. RSV-Luc was diluted to 2.55×10^4^ TCID50 per well and incubated with serial dilutions of monkey sera or BAL samples for 1 h at 37°C. Next, the serum-virus mix was applied to HEp2- cells, which had been seeded at 1×10^4^ cells/well the day before. Four hours after incubation at 37°C, 2% DMEM was added to the cells (2% FCS, 2 mM L-Glutamine, 100 units/ml penicillin/streptomycin). After 24 h at 37°C, cells were lysed with Glo Lysis Buffer 1x (Promega) for 15 min at 37°C. Luciferase luminescence was detected after 3 min incubation with Bright-Glo™ Luciferase Assay System (Promega). The plates were acquired on a microplate luminometer (VICTOR X5, PerkinElmer) using PerkinElmer 2030 Manager Software. The 50% plaque reduction neutralization titers (PRNT50) were defined as the highest dilution that inhibited more than 50% of plaques observed in eight infected control wells without serum treatment.

### qRT-PCR for viral load detection

Viral RNA was isolated from nasal and oropharyngeal swabs or BALF using the NucleoSpin® RNA Virus Kit (Macherey-Nagel) according to the manufactureŕs instruction. The quantity of viral RNA copies was determined by reverse transcription quantitative PCR (RT-qPCR) using GoTaq® 1-Step RT-qPCR System Kit (Promega). The following primers were used to amplify a sequence from the RSV nucleoprotein (for: AGATCAACTTCTGTCATCCAGCAA; rev: GCACATCATAATTAGGAGTATCAAT). In vitro transcribed RNAs were used as standards and the detection limit was 5 copies per reaction, which corresponds to 360 copies/ml.

### T cell ELISpot

IFN-y ELISpot assay was performed using commercially available reagents (Mabtech), as previously reported^36^. Briefly, triplicates of 1 × 10^5^ fresh peripheral blood mononuclear cells (PBMC) per well were seeded on 96-well plates (MultiscreenHTS IP filter plates; Millipore; #MSIPS 4510). Cells were either stimulated with 15-mer peptide pools covering the RSV F-protein (2 µg/ml of each peptide); with SEB as a positive control (Sigma # S4884-1mg; final concentration: 50ng/ml), with peptide pools of HCV (GeneScript; final concentration: 2µg/ml) as a negative control or left unstimulated. Spot numbers were analyzed on an ELISPOT reader (BIOSYS Bioreader 6000V)

## Results

In this confirmatory, dual-center NHP study, the immunogenicity and efficacy of a heterologous DNA prime – adenoviral vector boost immunization against RSV was analyzed in rhesus macaques. The study followed an adapted protocol from a previous exploratory study^29^ and was performed at two independent primate centers to increase internal and external validity. We had three experimental treatment groups referred to as RSV, Mock and comparator group (cohort characteristics, see table 1). The first two groups comprised the vaccine group (RSV) and the respective control group (Mock), which both received an initial DNA prime immunization applied via intramuscular electroporation at week 0. The antigens encoded by the plasmids were codon-optimized versions of either the RSV-F protein or the IAV-derived proteins hemagglutinin and nucleoprotein as unrelated Mock antigens. The same antigens were encoded by the Ad5-based booster vaccines, which were applied via the MAD720 device as an oropharyngeal spray at weeks 8 and 28 (Fig.1). Since natural immunity acquired by primary RSV infection does not provide long-term protection against RSV re-infection, we aimed at directly comparing natural immunity to our vaccine-induced immunity. Therefore, the comparator group was primarily infected with RSV at the time point of our first mucosal boost immunization at week 8. Since these animals served also as a comparator group in a subsequent IAV vaccine study, which is not part of this manuscript and will be reported elsewhere, the rhesus macaques received at week 24 and 28 two doses of a quadrivalent inactivated influenza vaccine, which should be irrelevant for RSV. To determine efficacy, all animals were challenged by RSV at week 40 and viral loads were measured in upper and lower respiratory tract samples.

### Primary RSV infection in the comparator group

The comparator group was included in the confirmatory study design to mimic the mucosal immune response in naturally RSV infected individuals. To confirm efficient infection of the upper and lower airway by our newly prepared RSV stock in RSV-naïve animals, nasal and throat swabs were collected on days 0,3 and 6 and a BAL was performed on day 6, which was around the peak of RSV replication in our previous exploratory study. Viral RNA was detectable in all but one animal (BPRC), but the strength of viral replication seemed to be slightly lower in BPRC-animals than in the ones housed at DPZ (Fig.2). In the latter, the inner-group variations of the replication kinetic was rather low and we detected between 10^6^-10^9^ copies of viral RNA/ml in the upper respiratory tract (Fig.2 A, B) and between 10^3^-10^6^ copies/ml in the BAL (Fig.2 C). In contrast, at the BPRC, two animals exhibited very low viral loads and seemed to be hardly infected, while the two others had similar replication kinetics to the ones at DPZ, but with overall 1-2 logs lower vRNA copy numbers in nasal and throat swabs (Fig.2 A, B). Since we observed substantial differences in our experimental data between the two centers, we will present and discuss all following results separately for each center.

**Fig. 2:**
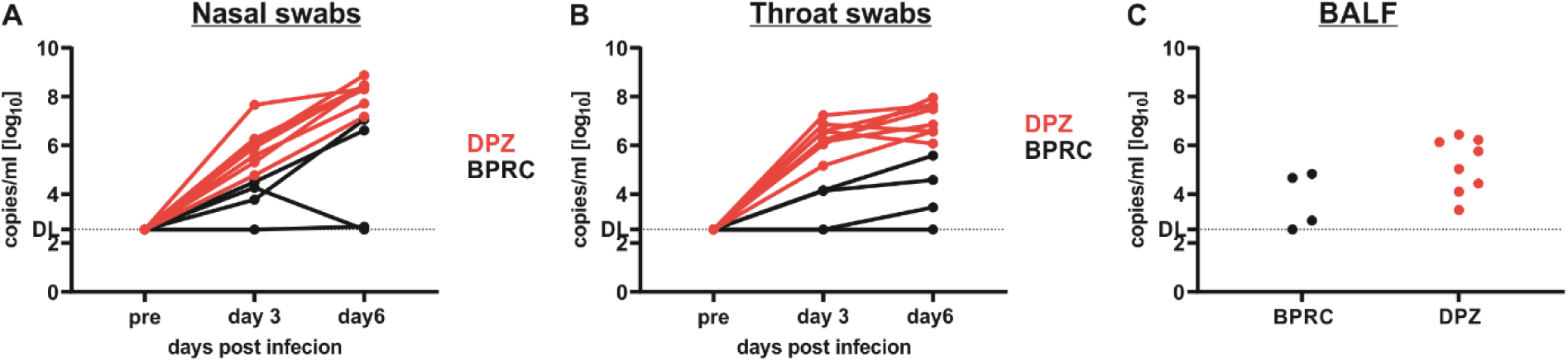
Viral loads after primary RSV infection in comparator group. Animals of the comparator group were infected with 1×10^5^ PFU macaque-adapted RSV at week 8. Nasal (A) and throat (B) swabs were collected pre-challenge and on day 3 and 6 post infection, while BALF (C) were collected on day 6. Viral loads were determined via qRT-PCR and copy numbers of vRNA/ml were indicated for each animal as individual curve. Red symbols and line represent animals from the DPZ, while black symbols and lines represent animals from BPRC.

### Humoral immune response

Initially, we analyzed longitudinally the antibody levels of RSV-F specific antibodies able to bind its target in the natural membrane-embedded conformation by our flow-cytometric assay. Here, we observed RSV-F specific IgG antibodies in sera (Fig.3 A, D) and BAL (Fig.3 B, E) as well as RSV-specific IgA antibodies in BAL (Fig.3 C, F). In all RSV-immunized animals, successful seroconversion was demonstrated by the presence of RSV-specific IgG in sera, but again the kinetics and the strength of the responses differed for the two centers (Fig.3 A, D). Interestingly, all animals at the DPZ developed anti-RSV-F IgG antibodies within 7 weeks after the DNA immunization, while only 1 out of 4 animals seroconverted at the BPRC during this period. However, the peak responses for F-specific serum IgG after the final vaccination were in the range of 30-70 µg/ml and thus in the same range for both centers (Fig.3 A, D). Additionally, the primary infection induced in all 12 animals of the comparator group detectable IgG levels, which were overall lower than in the RSV-immunized animals and mirrored the degree of RSV replication (Fig.2). The two animals with very low viral loads at the BPRC also had only marginal levels of F-specific IgG in the serum (Fig.3 D). Unexpectedly, animals of the Mock-treated group at the DPZ seroconverted for F-specific IgG between week 10 and 12, while the Mock-treated animals at the BPRC remained negative for RSV-specific antibodies throughout the immunization phase. In the DPZ Mock-group, the increase of F-specific IgG was delayed by two weeks in comparison to the comparator group, but reached similar peak levels, which might indicate an unintended cross-infection within the housing room.

**Fig. 3:**
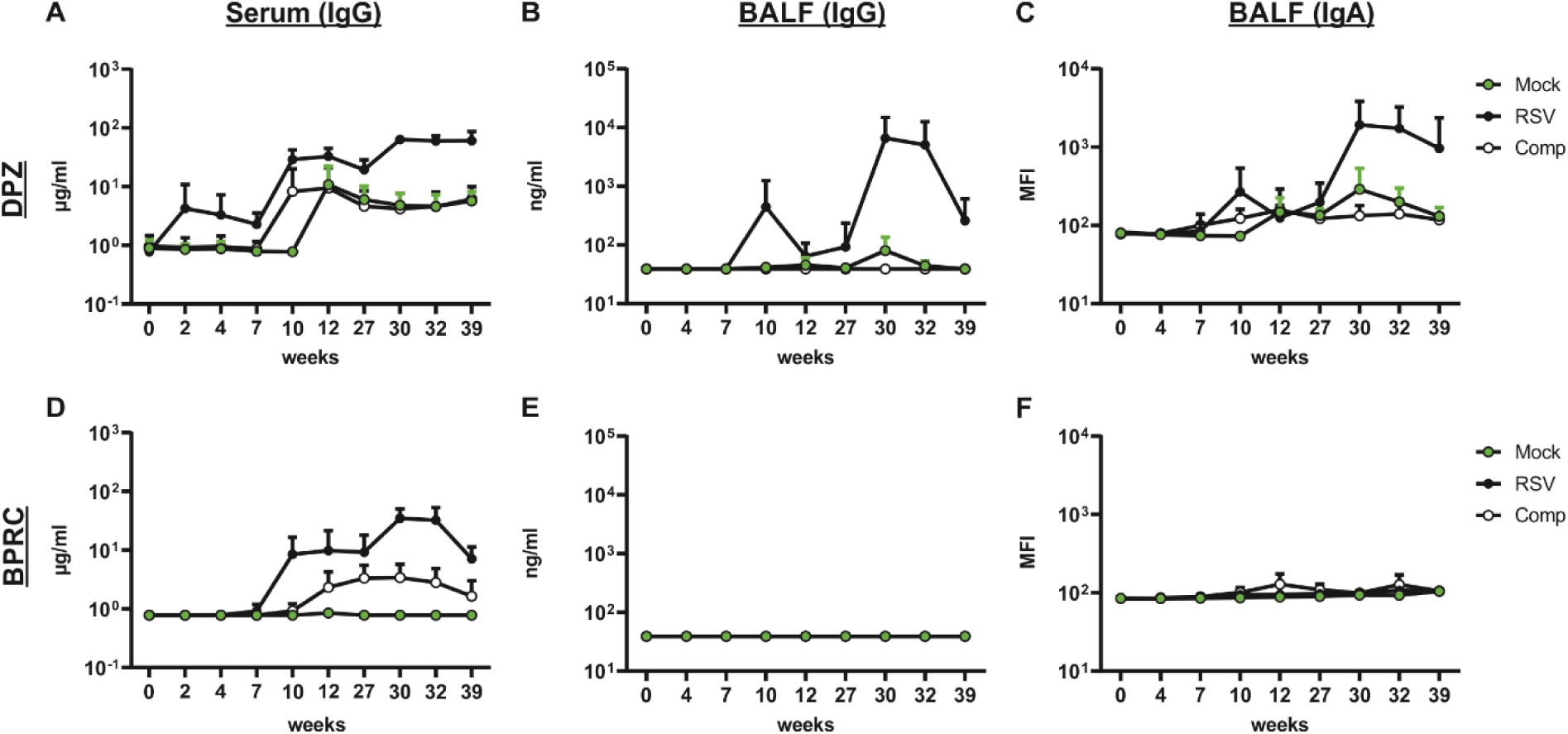
RSV-F specific antibody responses. Animals were immunized as illustrated in Fig.1. Serum (A,D) and BALF (B,C,E,F) were collected at indicated time points. RSV-F specific IgG (A,B,D,E) and IgA (C,F) antibodies were determined by a flow-cytometric antibody binding assay using stably RSV-F expressing cells as target. The amount of IgG in serum and BAL was calculated according to a standard curve with the monoclonal antibody palivizumab and shown as µg/ml. For IgA, the mean fluorescence intensity is indicated. The symbols represent the mean and the error bars the SEM of each group (n=8 for DPZ; n=4 for BPRC)

Interestingly, the primary RSV infection failed to induce substantial mucosal antibody responses (Fig.3 B, E). In sharp contrast, our heterologous vaccination regimen led to detectable F-specific IgG and IgA in the BALF of all eight DPZ-animals (Fig.3 B, C). The level of IgA and IgG in BAL peaked two weeks after the second mucosal Ad5 booster and then gradually declined until the time of challenge (W39). Surprisingly, we could not detect any F-specific antibodies in the BAL of BPRC animals (Fig.3 E, F).

In addition to analysis of RSV-F binding antibodies, we measured the neutralization capacity of the systemic and mucosal antibodies using a recombinant, luciferase-expressing RSV reporter. In line with the flow-cytometric assay, the highest neutralization titers were observed in the RSV vaccine group from DPZ after the second Ad5 booster with about 3-fold higher titers than in the respective BPRC group (Fig.4 A, C). In the comparator and mock-treated groups, modest neutralization could be detected only in a small fraction of the animals.

**Fig. 4:**
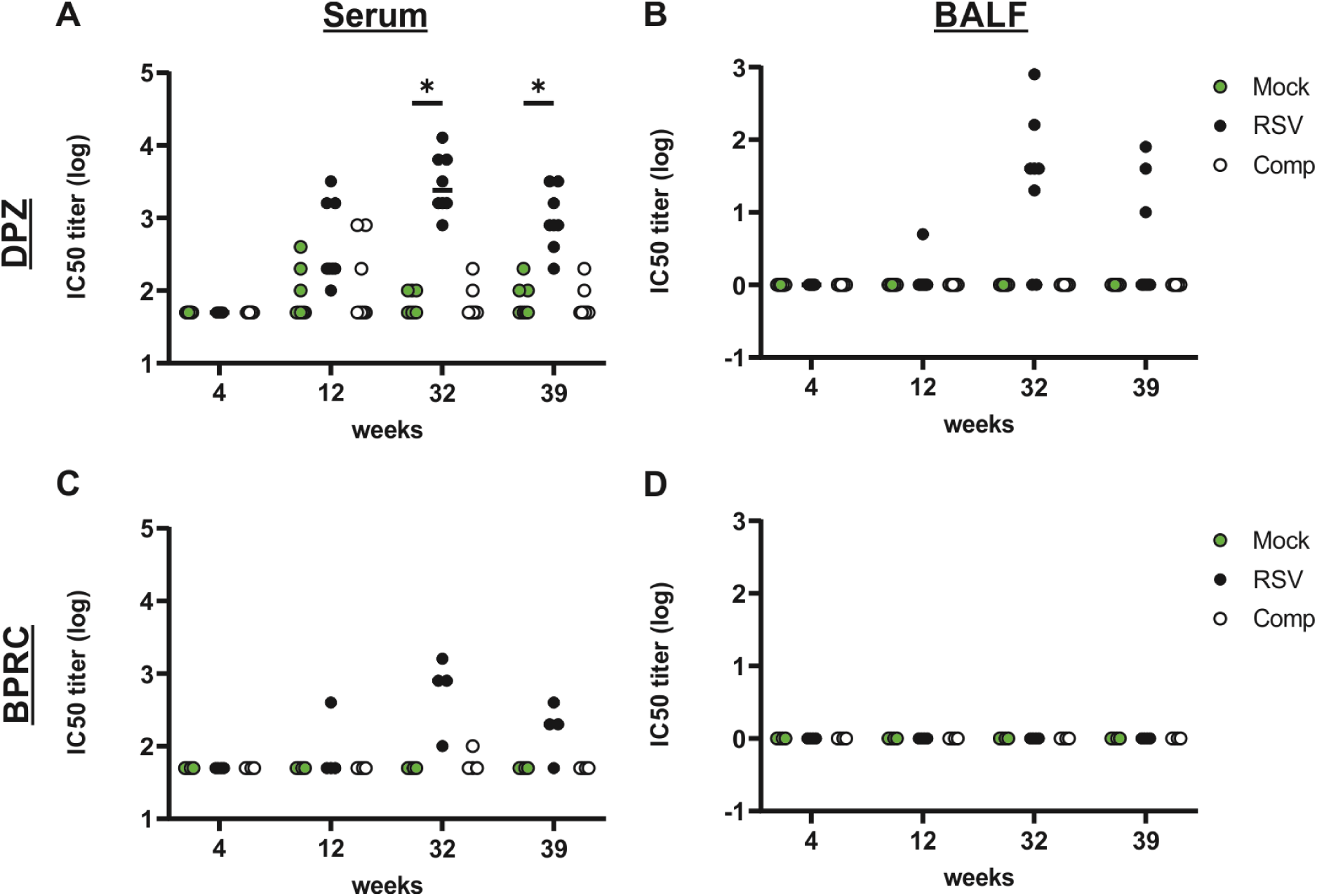
Neutralizing antibodies. Animals were immunized as illustrated in Fig.1. Serum (A,C) and BALF (B,D) were collected at indicated time points. Neutralizing antibodies were analyzed using a recombinant luciferase-expressing RSV reporter. The reciprocal value of the sample dilution preventing 50% of the infection events is presented as IC50 tier, which is indicated for each individual animal. (n=8 for DPZ (A,B) ; n=4 for BPRC (C,D)) Statistical significances were analyzed by two-way ANOVA followed by Dunnett’s multiple comparisons test and p values indicate significant differences (*p < 0.05).

Again, neutralizing antibodies in the mucosal compartment was detected only in the DPZ animals and not in the ones housed at BPRC (Fig.4 B, D). However, at week 39 only three animals had still IC50 titers above 10, which indicates the lower sensitivity for BAL samples, which might be explained by the substantial dilution of locally present antibodies due to the large BAL volume (Fig.4 B).

### Cellular immune response

RSV-specific T-cell responses were evaluated by functional ELISpot assay and ICS after in vitro restimulation with overlapping peptide pools spanning the whole F protein. Unfortunately, the frozen PBMC samples showed a strong reduction in their responsiveness in our IFN-γ ELISpot after thawing compared to the freshly isolated PBMCs and gave inconclusive results with considerable drop-outs. Therefore, we present only data from freshly processed PBMC samples, which were exclusively available for the DPZ animals. However, these 24 animals gave insights in the kinetics of the systemic T-cell response and allowed statistical analyses. Here, the DNA immunization already primed a robust T-cell response detectable at week 4 in all RSV- vaccinated macaques (Fig.5). Interestingly, in the same animals only low number of IFN-γ producers could be detected among the PBMC two weeks after the first mucosal booster immunization. In contrast, the second mucosal booster resulted in strong recall responses and most animals reached their maximum number of IFN-γ spots, which then gradually declined again until week 39. Very limited and only temporary IFN-γ responses were detected in the Mock-treated animals over the whole observation period. In the comparator group, the primary RSV infection at week 8 induced a robust, but short-lived systemic T-cell responses in all eight animals. However, neither the Mock nor the comparator animals had detectable levels of IFN-γ producing T-cells one week before the challenge virus exposure took place (Fig.5).

**Fig. 5:**
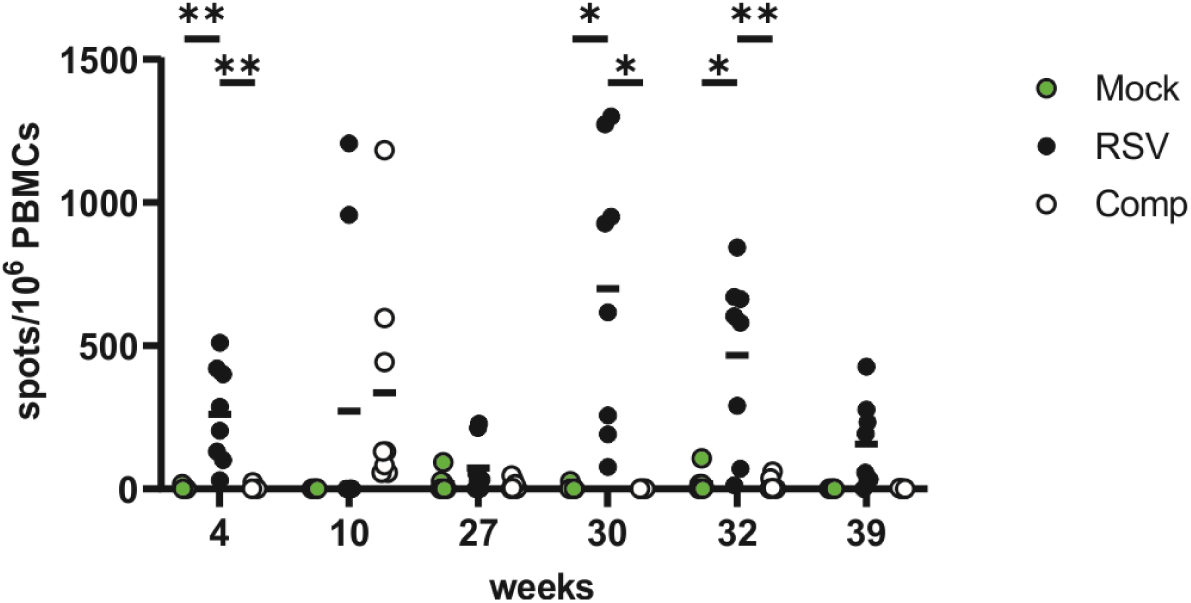
RSV-F specific systemic T-cell response. Animals were immunized as illustrated in Fig.1. PBMCs were freshly isolated from the animals at DPZ and restimulated with RSV-F specific peptide pools or respectice controls. RSV-F-reactive T-cells were analyzed by IFN-ELISpot. Each symbol represents an individual animal (n=8 for DPZ). Statistical significances were analyzed by two-way ANOVA followed by Dunnett’s multiple comparisons test and p values indicate significant differences (*p < 0.05, **p < 0.01).

In addition to the systemic T-cell response, antigen-specific CD4 and CD8 T-cells were analyzed in the BAL by intracellular cytokine staining to evaluate the mucosal responses (Fig.6, gating in Suppl. Fig.1). In accordance with the ELISpot results, the peak response was observed in the RSV group two weeks after the second booster immunization (W30) and was characterized by high rates of cytokine-producing CD4 (Fig.6 A-D) and CD8 T-cells (Fig.6 E-H). Moreover, we identified TRM subpopulations within the total CD4 and CD8 T-cell subsets by using the markers CD69 and CD103 (Suppl.Fig.1). In almost all RSV vaccinees from both centers, we were able to detect IFN-γ producing CD4 TRM in the BAL at varying frequencies ranging from 1% to 23% of all CD4 TRM cells (Fig.6 B, D). Two weeks after the primary RSV infection (W10), antigen-specific CD4 and CD4 TRM cells were also detectable in the animals of the comparator group, but declined then gradually until the challenge (W39). Except for some sporadic cytokine responses in individual animals, we did not detect RSV-specific T-cell responses in the mock group. Overall, the CD8 T-cell responses followed the same trends and kinetics as observed for CD4 T-cells, but were less frequently measurable than antigen-specific CD4 T-cells (Fig.6 E-H). This might be attributable to the characteristics of the peptide pools used for re-stimulation. Those consisted of 15-mers and might be not optimal for loading of MHC-I molecules. Again, the peak responses for CD8 and CD8 TRM were observed in the RSV-immunized animals after the second booster (week 30/32).

**Fig. 6:**
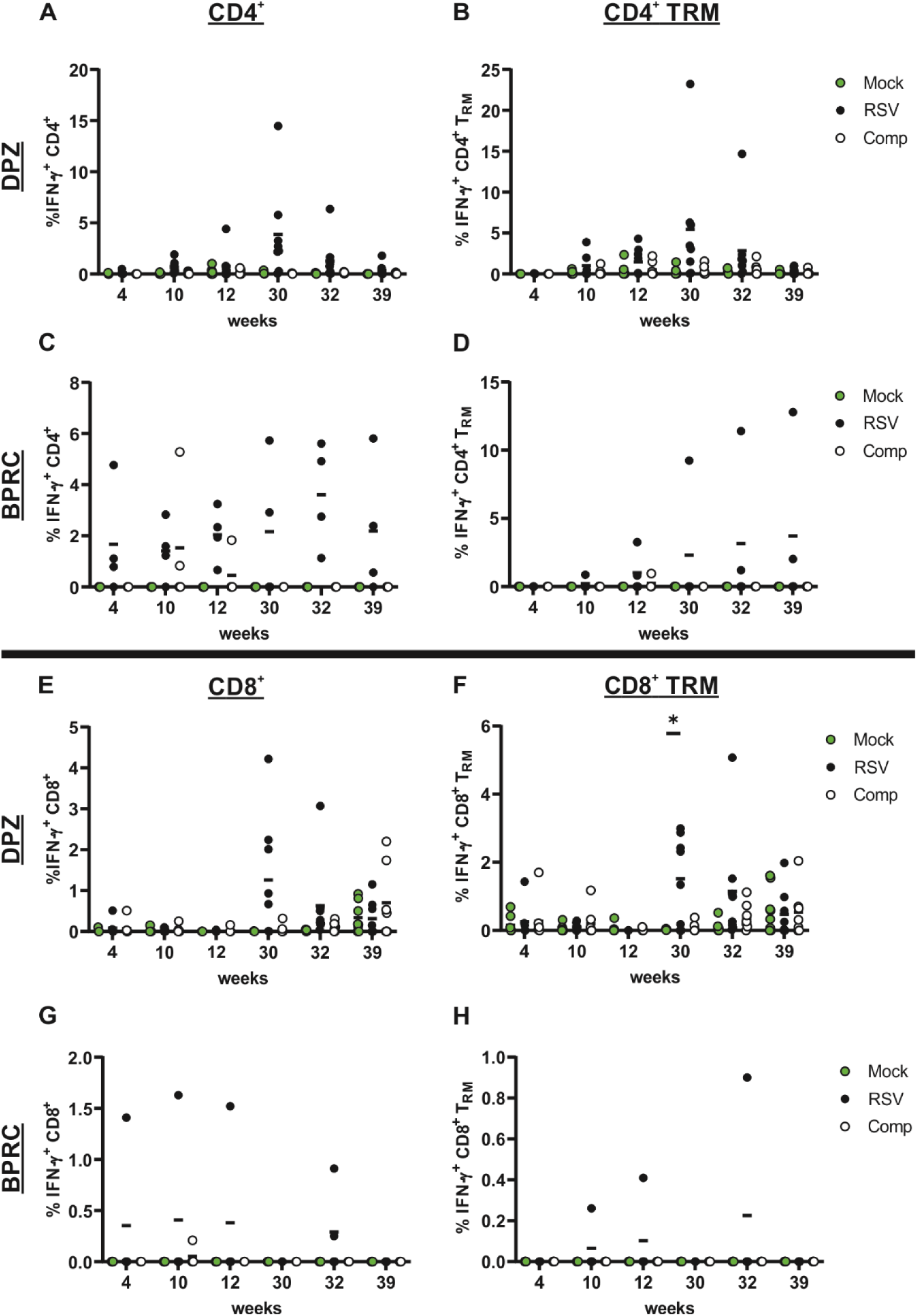
RSV-F specific mucosal T-cell response. Animals were immunized as illustrated in Fig.1. BAL cells were freshly isolated from the animals and restimulated with RSV-F specific peptide pools or respective controls. RSV-F-reactive T-cells were analyzed by intracellular cytokine staining. The percentage of IFN-γ^+^ CD4^+^ T cells (A,C), CD4^+^ TRM (B,D), CD8^+^ T cells (E,G) and CD8^+^ TRM (F,I) is indicated for each animal from both centers. Each symbol represents an individual animal (n=8 for DPZ; n=4 for BPRC). Statistical significances were analyzed by two-way ANOVA followed by Dunnett’s multiple comparisons test and p values indicate significant differences (*p < 0.05, **p < 0.01).

### Vaccine efficacy

The primary end-point of this study was the confirmation of vaccine efficacy in an RSV challenge experiment. Thus, all animals were exposed to RSV at week 40. Since RSV infection does not induce any respiratory symptoms in rhesus macaques, the viral loads in the upper and lower respiratory tract were monitored by qRT-PCR as already described for the primary infection in the comparator group. Again, we observed striking differences in the infection kinetics between the two centers (Fig. 7 A-I). At the BPRC, all mock treated animals had high copy numbers of viral RNA in the nasal as well as in the throat samples between days 3-9 post infection before gradually clearing viral infection (Fig.7 D, E). The RSV vaccinated animals were almost completely protected against the upper respiratory tract infection with only two nasal samples in which RNA copy numbers were slightly above the detection limit on day 3. In the comparator group, natural immunity induced by the primary RSV infection was also partially protective. Here, only in one out of four animals a typical viral RNA replication kinetic could be observed in the upper airways, but also at significantly lower levels than in the mock group (Fig.7 D, E). The degree of protection induced by the primary infection and our RSV vaccination was similar in the DPZ animals (Fig.7 A, B). Here, seven out of eight animals in the comparator group had detectable levels of vRNA in the nose on day 3 post infection and three animals were repeatedly RNA positive over the course of infection. In the RSV group, four and six animals stayed completely negative for vRNA in the nose and the throat, respectively. The few vRNA positive samples exhibited viral loads just above the detection limit. Unexpectedly, the mock group from DPZ revealed considerably lower vRNA levels in the upper airway (Fig.7 A, B) than those observed in the DPZ comparator group after primary RSV infection (Fig. 2) and those in the mock group from BPRC (Fig.7 D, E). Kinetics and height resembled those of the comparator group after its secondary infection, which is in line with the previously described anti-F immunity detected in these animals (Fig.3-5). Considering the total RNA level (area under the curve) for all 12 animals of the groups, there was a significant difference between the Mock- and RSV-vaccinated animals for both upper respiratory tract compartments (Fig.7 G, H). However, the viral load levels differed substantially depending on the nonhuman primate center performing the study. Infection of the lower airways was overall very limited. We found very low copy numbers of viral RNA in all four mock-immunized animals from the BPRC, but in none of the other 32 animals (Fig.7 C, F, I).

**Fig. 7:**
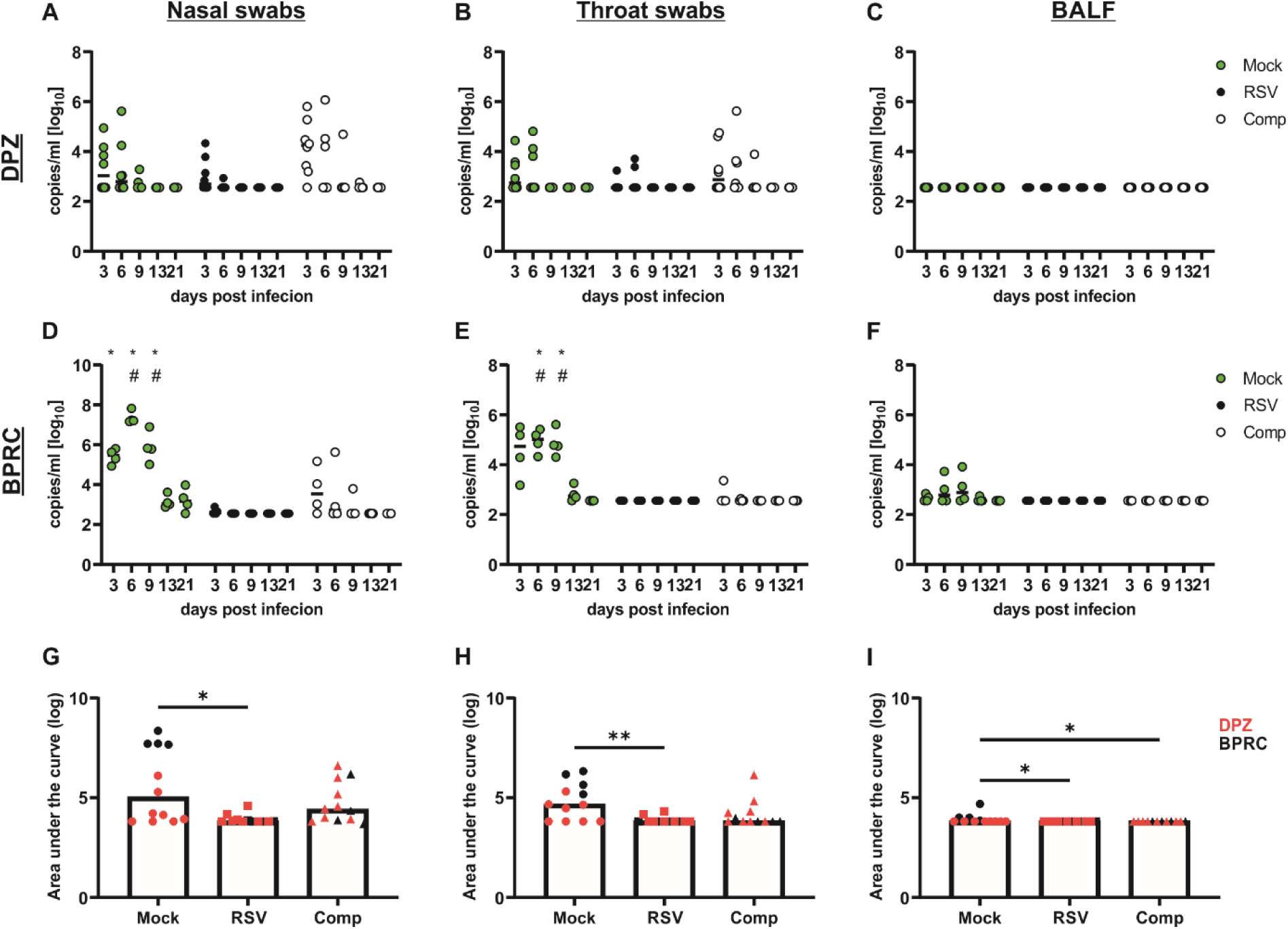
Viral loads after RSV challenge. Animals were immunized as illustrated in Fig.1 and infected at week 40 with 1×10^5^ PFU macaque-adapted RSV. Nasal (A,D) and throat (B,E) swabs and BALF (C,F) were collected at the indicated time points. Viral loads were determined via qRT-PCR and copy numbers of vRNA/ml were indicated for each animal at both centers (A-F). Statistical significances were analyzed by two-way ANOVA followed by Dunnett’s multiple comparisons test and p values indicate significant differences (*p < 0.05, **p < 0.01). (G-I) To estimate the viral load over the whole observation period, the area under the curve was calculated for each animal. Here animals of both centers were shown in a combined figure. Red symbols represent animals from the DPZ, while black symbols represent animals from BPRC. The Bars represent the median of the respective group. (n=8 for DPZ; n=4 for BPRC). Statistical significances were analyzed by Kruskal-Wallis test followed by Dunn’s multiple comparisons test (*p < 0.05, **p < 0.01).

In addition to the viral RNA measurement, post-challenge antibody levels were evaluated as indirect measure of viral replication (Fig. 8). Almost no increase in RSV-F specific antibody levels was detected in RSV-immunized animals from DPZ. By contrast, the comparator and mock group had stronger anamnestic responses suggesting less viral blocking and higher antigen exposure during the challenge (Fig.8 A, B). The Mock treated animals at DPZ showed again very similar response kinetics as the comparator group. The increase in RSV F-binding antibodies was mirrored by increased neutralization capacity of serum from these animals (Fig.8 C). At BPRC, F-specific antibodies became also detectable in Mock-treated animals after the challenge, albeit remained at lower levels than in the two other groups with almost no neutralizing activity (Fig.8 D-F), which resembles the antibody response after the primary RSV infection in the comparator group (Fig. 3, W12). The rather limited booster response in the RSV-immunized animals might be indicative of an early restriction of viral replication by local immune players, such as IgA and/or F-specific TRM.

**Fig. 8:**
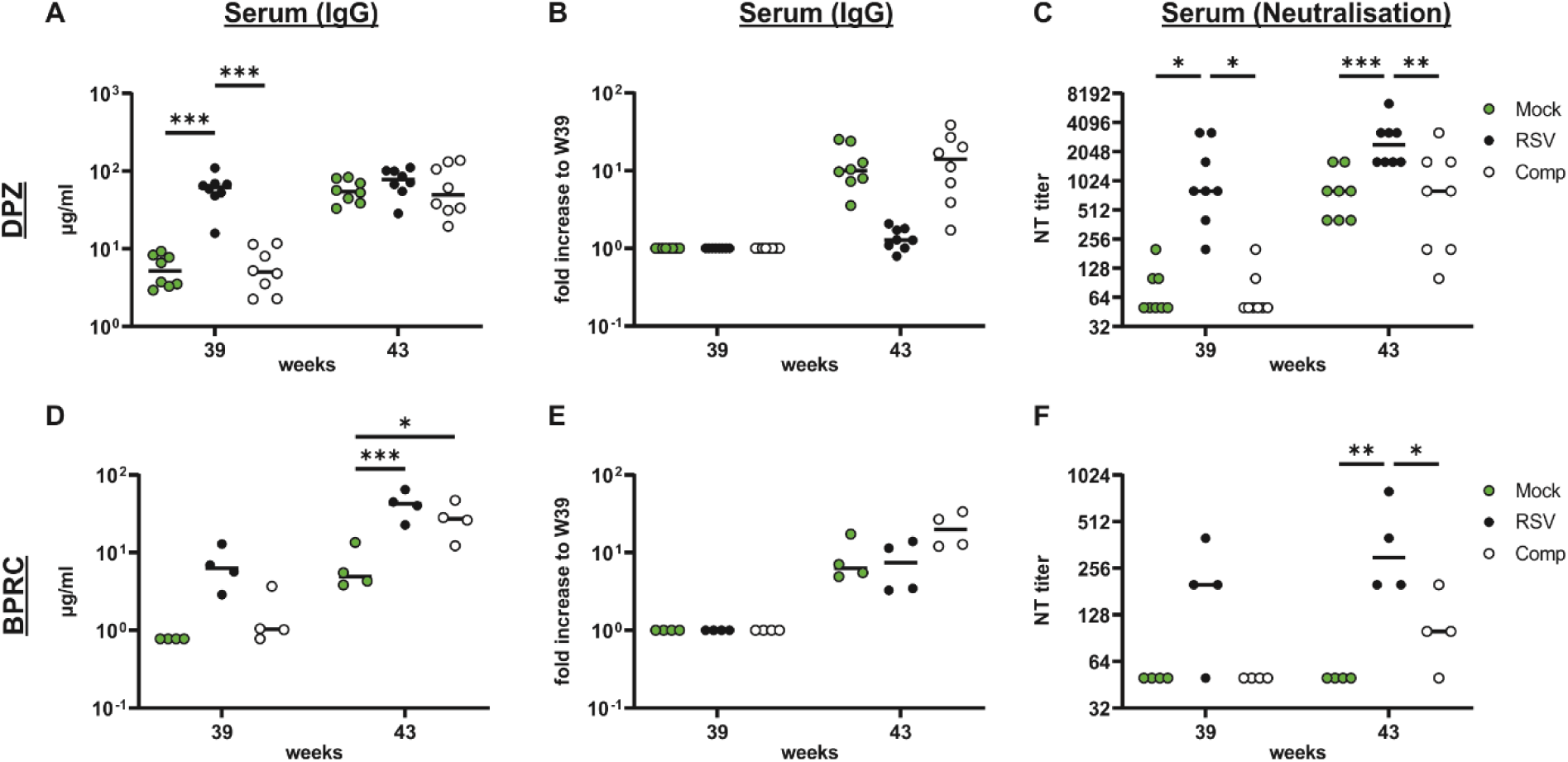
Anamnestic antibody response after RSV challenge. Animals were immunized as illustrated in Fig.1 and infected at week 40 with 1×10^5^ PFU macaque-adapted RSV. RSV-F specific antibody responses were determined pre (W39) and post (W43) RSV challenge via flow cytometric antibody binding (A,B,D,E) and neutralization assay (C,F). The total amount of IgG (A,D) as well as the fold increase between pre/post challenge (B,E) are indicated for each individual animal from both centers. The IC50 NT values were calculated as described and also indicated for each individual animal (C,F) (n=8 for DPZ; n=4 for BPRC). Statistical significances were analyzed by two-way ANOVA followed by Dunnett’s multiple comparisons test and p values indicate significant differences (*p < 0.05, **p < 0.01).

In summary, these dual-centric NHP study generated further evidence that our heterologous prime-boost regimen is highly immunogenic, capable of inducing mucosal immunity and protects from subsequent RSV infection. Of note is the high degree of variation between the two primate centers and the unexpected outcome of the DPZ Mock group.

## Discussion

Here, we report on a multi-centric NHP study planned as a confirmatory study for our previous exploratory study on a new vaccination strategy for prevention of respiratory tract infections with RSV and the difficulties faced during the experimental process. Indeed, there are overall and specifically in the field of immunology or infectious disease research very few reports on multi-centric NHP studies, which already indicates the particular character of our study. One study evaluated the safety and immunogenicity of a licensed, attenuated measles vaccine at different primate centers, but the analyses was primarily focused on seroconversion rates^37^. However, no complex vaccination regimen with preclinical vaccine candidates have been reported in such a multi-centric NHP approach so far.

Our vaccination schedule consists of a primary, intramuscular DNA vaccine applied by electroporation followed by two oropharyngeal spray immunizations with replication-deficient rAdV encoding for a codon-optimized, non-stabilized RSV-F protein^29^. To increase external validity of our exploratory results, we included a comparator group that was exposed to RSV to induce natural immunity to infection. This usually does not confer long-term protection against re-infection in humans^5–7^. The primary RSV infection in rhesus macaques induces no or only mild clinical symptoms, but results in robust viral replication in the upper and lower respiratory tract when the virus is installed via the intranasal route^29,34^. In the comparator groups, we could confirm those replication patterns, but observed lower viral loads in the lower respiratory tract compared to previous experiments. This slight difference in viral infectivity might be a result of batch-to-batch variations during RSV stock preparation. Here, a newly raised stock of macaque-adapted RSV^34^ propagated on TMK cells was used in this study. In addition, the infection of the upper respiratory tract differed in strength for the two centers as viral loads were consistently lower in the animals at BPRC. In light of the observed differences in the mucosal antibody responses (discussed below), we identified minor differences in the anesthesia procedure between the two centres during the phases of primary RSV infection and the oropharyngeal spray immunization. While a combination of ketamine, xylazine, and atropine was used at the DPZ as in our previous study, the standard procedure at BPRC consists of ketamine mixed with medetomidine followed by later application of atipamezole as antagonist for faster recovery. This difference in depth and length of sedation might have influenced the duration of exposure of the mucosal surface to the challenge virus and the oral vaccines and thereby subsequently the strength of viral replication or antigen expression. Deeper sedation periods can more efficiently suppress sneezing or coughing reflexes and result in deeper and slower breathing. In a murine influenza A virus infection model, more severe disease progression and a higher degree of viral replication in the lower respiratory tract was observed upon intranasal infection under deep xylazine/ketamine anesthesia compared to light isoflurane inhalation^38^. Similar findings have been reported for bacterial pneumonia induced by Streptococcus pneumoniae ^39,40^. Furthermore, other physiological or immunomodulatory alterations induced by the different anesthesia protocol (reviewed in ^41^) could not be excluded as potential co-factors for the differential outcomes for the two centers.

As described above, the two different anesthesia protocols also were used for the oropharyngeal spray immunizations and might partially explain the lower anti-F antibody levels seen in the BAL samples of the BPRC animals. While anti-F IgG and IgA antibodies were readily detectable in the BAL of DPZ animals after the first AdV spray application and reached peak levels two weeks after the second boost, anti-F antibodies were nearly absent in the BAL and only serum IgG could be detected at BPRC. In a recent study, Tang et al. reported efficient induction of mucosal IgA responses in mice only when the AdV was delivered in large volumes reaching the lower respiratory tract, but not when smaller volumes were applied^42^. Similarly, the AdV vaccination under the mild anesthesia protocol at BPRC might have restricted the transgene expression more to the upper respiratory tract. Unfortunately, we do not have samples from the nose to analyze IgA responses in the upper respiratory tract, which is clearly a limitation of our study.

Overall, the humoral response seemed to be less potently induced in the BPRC animals compared to the DPZ ones. This became already obvious after the DNA electroporation. At DPZ, serum IgG appeared in response to the DNA treatment already after two weeks and increased with each booster immunization reaching a stable plateau until the challenge thus confirming the data from our initial NHP study ^29^. Although we used the identical DNA vaccine preparation and electroporation device at the BPRC, only one animal seroconverted after DNA administration. To avoid biases in the analyses, we investigated the humoral responses in samples from both centers side-by-side in the same laboratory to minimize technical variations. Therefore, other non-obvious co-founders, such as center-specific diets, animaĺs microbiota or experimental history, might have also influenced the immunological outcome of our vaccination regimen. In summary, RSV F-specific antibodies were induced in all DNA-AdV vaccinated animals, but the absolute magnitude was lower at the BPRC. This was also confirmed on a functional level by the neutralization assays.

Interestingly, the mucosal T-cell responses were more comparable between the two centers. In line with several other studies from our and other labs, a systemic prime mucosal boost immunization induced substantial amounts of antigen-specific CD4 and CD8 TRM ^21,29,43–46^. Due to the use of 15-mer peptide pools and potentially less efficient presentation on MHC I ^47^, the CD8 response might be even underestimated and therefore was not as consistently detected as the CD4 T-cell response. In addition, RSV F-specific T-cells in the PBMC compartment were identified by IFN-γ ELISpot analyses. These cells were already induced by the DNA electroporation and boosted by the oropharyngeal AdV spray application confirming our data from the exploratory studies ^29^.

In our comparator group, natural immunity induced by a primary RSV infection was characterized by F-specific IgG with moderate neutralization capacity in serum and RSV-F specific CD4 T-cell responses in BAL and PBMCs. However, mucosal antibodies and CD8 TRM responses were almost absent after the primary infection. This might be in sharp contrast to the human situation, where local immunity at the mucosal site is reported in form of IgA and TRM partially protective against re-infection^48,49^. It represents one of the general limitations of RSV infection in the semi-permissive rhesus macaque model. Nevertheless, we observed considerable reduction in viral load of the upper airways in animals from both centers after the RSV challenge infection. Surprisingly, the Mock control group at DPZ developed immune responses comparable to those of the comparator group and demonstrated a similar level of protection following the RSV challenge. Due to welfare reasons animals of the comparator group could not be housed separately from the other two groups suggesting a potential cross-transmission within the animal facility. However, horizontal RSV transmission among rhesus macaques has not been reported so far and is questionable in the light of the semi-permissive nature of this RSV model. Similarly, a cross-species transmission from an infected animal care taker to the macaques would be principally possible as such human-to-monkey transmission have been reported for human metapneumovirus in a colony of wild chimpanzees^50^. Nevertheless, due to the strict safety measures in the animal facilities, i.e. wearing of FFP2 face masks, clothing changes etc., this is also not a very likely scenario. Furthermore, we could exclude any contamination of the respective adenoviral vector preparations used for vaccination. We transduced A549 cells with aliquots from the vaccine preparations used for the immunization as well with the initial AdV stocks and confirmed the correct antigen expression for each vector. An additional argument against an underlying contamination as reason is the fact, that the vaccine doses for both centers were generated simultaneously and we did not observe the same kind of RSV-F reactive response in the Mock group at BPRC.

To summarize, we cannot conclusively answer the question of why animals of the DPZ Mock group developed RSV-specific humoral responses following RSV infection of the DPZ comparator group, particularly as vertical transmission cannot be excluded. The relatively low copy numbers of vRNA in the mock group after RSV challenge were the reason why we could not confirm the efficacy of our experimental vaccine at the DPZ. By contrast, high efficacy was observed in the RSV group of BPRC preventing viral replication in the upper respiratory tract. However, RSV replication was almost completely absent in all animals having received the heterologous prime-boost immunization independent of the housing location. Systemic neutralizing antibodies in combinations with rapid reactivation of F-specific TRM might have limited the degree of viral replication at the mucosal entry site, which have been described as correlates of protection before ^48,49,51^. The contribution of mucosal IgA antibodies in this study is difficult to assess, because we were unable to detect F-specific antibodies in the BAL of animals housed at BPRC. However, we did not analyze other mucosal samples, such as nasal or throat swabs, and could thereby not exclude that IgA might be present at the viral entry sites of the upper respiratory tract.

In the SARS-CoV-2 macaque model, an oropharyngeal spray immunization with AdV mounted SARS-CoV-2 specific IgA antibodies in serum, nasal washes and BALF in mRNA primed animals. These responses negatively correlated with viral RNA after SARS-CoV-2 challenge infection^43^. This is in line with several other studies demonstrating efficacy of heterologous systemic prime-mucosal boost strategies against respiratory viruses, such as SARS-CoV-2, Influenza A virus or RSV ^21,29,43–46^.

Collectively, our confirmatory multi-centric NHP study failed to reach the primary endpoint due to some unforeseen obstacles during the experimental phase. However, this might inform the community and generate awareness for the design of future multi-centric animal studies. Finally, our study still generated evidence for the potential of improved mucosal vaccines against emerging or re-emerging respiratory pathogens, which could be exploited to improve pandemic preparedness.

## Data availability

All data are included in the Supplementary Information or available from the authors, as are unique reagents used in this article. The raw numbers of all graphs are available in the Source Data File linked to this article.

## Acknowledgement

We thank Silke Schroedel and Christian Thirion from SIRION Biotech for providing the Ad5-based vector vaccines. Furthermore, we are grateful for the support of Rik de Swart (Department of Virology, Wageningen Bioveterinary Research, Lelystad, The Netherlands), who provided the macaque-adapted RSV strain. We thank Philipp Brytzki, Nicole Leuchte and Sandra Heine (all DPZ) for excellent technical assistance. This work was funded by the German Ministry for Education and Research (BMBF) under the project title RSV-PROTECT (AZ:. 01KC2007)

## Author contribution

MT, CSH, and KÜ conceived and designed the study. GK, PM, BR, PI, DL, IK, RV and SAMR, performed the experiments and collected the data. SM and OG performed statistical analyses and sample size calculations for the experimental setup. GK, WMB, EJR, NKV and provided critical methodology and reagents. MT, GK, PM, BR, RV, SM, CSH, and KÜ analyzed and interpreted the data. MT, CSH, OG and K.Ü obtained the funding. MT, GK, CSH and KÜ drafted the manuscript, which was then critically reviewed and approved by all co-authors.

## Notes

### Competing Interest Statement

The authors have declared no competing interest.

