## Supplementary Material for "A confirmatory, dual-centric non-human primate study on the efficacy of novel oropharyngeal spray immunization with an adenoviral vector vaccine against RSV – Important lessons learned"

### Supplementary Figure 1

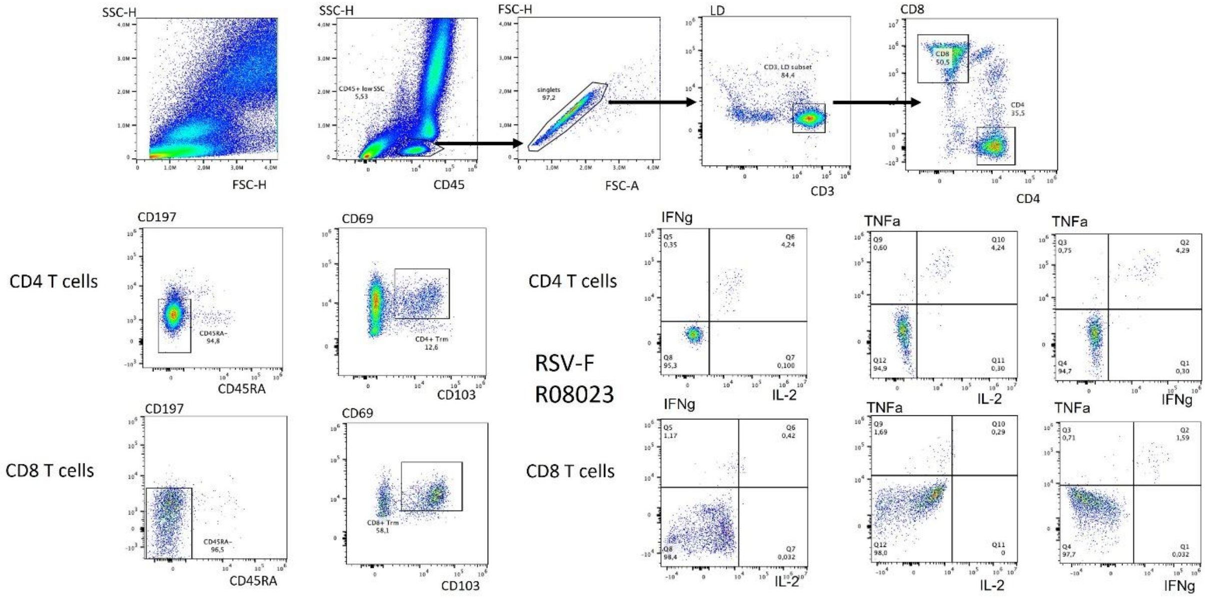


**Suppl.Fig.1: Gating strategy for intracellular cytokine staining of BAL cells**
Representative dot plots are shown for one BAL sample from the RSV vaccine group at BPRC (animal: R08023) which was stimulated with the FSV-F specific peptide pool.

### Supplementary Figure 2

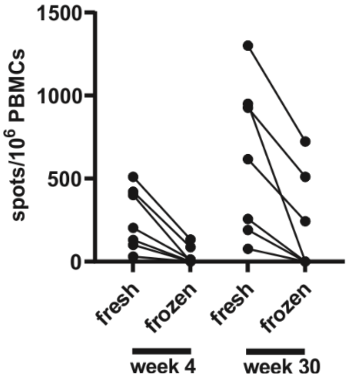


**Suppl.Fig.2: Comparison of ELISPOT analyses of fresh or frozen PBMC samples**
PBMCs were isolated at the indicated time points and either frozen and stored at -80° until the ELISPOT assays were performed or were directly restimulated for the ELISPOT assay. Samples from the RSV-immunized animals from DPZ were shown for the two time points (W4 and W30). Pairing of the individual samples are indicated by the lines.
